# Genetic Diversity and Structure of *Oryza rufipogon* Griff. Populations in the Philippines

**DOI:** 10.1101/2020.07.17.194829

**Authors:** Sandy Jan E. Labarosa, Neah Rosandra Sevilla, Dindo Agustin A. Tabanao, Nenita B. Baldo, Helen L.V. Ebuña, Joy M. Jamago

**Affiliations:** Department of Agronomy and Plant Breeding, College of Agriculture, Central Mindanao University, Musuan, Maramag, Bukidnon, 8710 Philippines; Plant Breeding and Biotechnology Division, Philippine Rice Research Institute, Science City of Muñoz, Nueva Ecija, 3119 Philippines

**Keywords:** *Oryza rufipogon*, wild rice, plant genetic resources, microsatellite, genetic diversity

## Abstract

*Oryza rufipogon* Griff. or ‘Rufi’ is the wild progenitor of the cultivated rice, *Oryza sativa* L. In the Philippines, Rufi was previously known to be found only in Lake Apo, Bukidnon. However, a new population was identified in Lake Napalit in the same province. Based on previous morphological diversity assessment, both populations are unique for at least three characters, i.e., leaf, culm, and awn lengths. Environmental parameters such as rainfall and air temperature also differed between the two lakes. With these, an assessment of Rufi’s genetic diversity at the molecular level is beneficial to further ascertain its usefulness in rice breeding and gain insights on its conservation status. Thus, this study estimated the degree of genetic diversity and determined the population structure of 41 samples of natural Rufi populations in the Philippines using SSR markers. A total of 98 genome wide polymorphic SSR markers were selected to examine the genetic diversity and structure of Rufi populations, along with seven rice cultivars for comparison. Results showed that Philippine Rufi populations have lower genetic diversity compared to cultivated rice accessions and other Rufi populations in Southeast Asia and China. This low genetic diversity suggested that Rufi populations might be in a genetic bottleneck, perhaps due to observed unsustainable farming practices near their habitat and lack of awareness of their importance. A significant population structure and differentiation were determined using the STRUCTURE and phylogenetic analyses. Population differentiation might be due to geographic isolation which prevented gene flow between the two populations and the unique climatic conditions between the two lakes.

## INTRODUCTION

*Oryza rufipogon* Griff. (nicknamed ‘Rufi’, 2n=2x=24) is the ancestor of the cultivated rice (*Oryza sativa* L.) (Gao et al., 2002; Song et al., 2005). For breeding, Rufi and 20 other wild rice species offer a reservoir of genetic diversity as essential sources of resistance and tolerance to some biotic and abiotic stresses, among others. Rufi has resistance mechanisms to various pests and diseases, such as bacterial leaf blight (Song et al., 2005), tungro (Song et al., 2005; Shibata et al., 2007), green leafhopper (Shibata et al, 2007), and brown planthopper (Li et al, 2010). It also has tolerance mechanism to several environmental stresses, such as aluminum toxicity (Nguyen et al., 2003), acid sulfate soils (Song et al., 2005; Bui and Nguyen, 2017), and cold temperature (Song et al., 2005).

Bon and Borromeo (2003) reported that the only natural populations of Rufi in the Philippines were in Lake Apo, Barangay Guinoyoran, Valencia City, Bukidnon. However, Jamago et al. (2012) recorded the existence of new natural populations in Lake Napalit, Barangay Pigtauranan, Pangantucan, Bukidnon. These new populations might harbor novel alleles useful for current and future rice breeding programs. Balos and Jamago (2013) compared some morpho-ecological parameters of Rufi populations in both lakes *in situ*. Their findings showed that Lake Apo populations (Apo) had longer leaves, culms, and awns than the Lake Napalit populations (Napalit). Also, rainfall was higher in Lake Napalit than in Lake Apo whereas, temperature was lower in Lake Napalit. These variations in morphological and ecological parameters between the two populations offer an opportunity for researchers and breeders to explore for their possible use in rice breeding programs.

This study estimated the magnitude of genetic diversity and determined the population structure of *in situ* populations of *O. rufipogon* Griff. in the Philippines using Simple Sequence Repeats (SSRs). Further, allelic patterns and degree of population differentiation across the populations were also calculated. Information on Rufi’s genetic diversity and population structure would provide more clarity on its potentials for utilization and even insights for its conservation.

## MATERIALS AND METHODS

### Leaf Sampling and Genotyping

Twenty-seven and 14 Rufi strains from Lake Napalit and Lake Apo populations, respectively, were sampled *in situ* (Figure 1) in 2014 spaced at least 30 meters to minimize the collection of leaves from possible clonal plants. Leaf sampling was done by cutting about 4-5 inches from the tips of flag leaves. Leaf cuttings per strain were placed in individual 200 ml centrifuge tubes that were immediately placed in an icebox before storing at –20° C until DNA extraction. Total gDNA was extracted from lyophilized leaf tissues following the modified cetyltrimethylammonium bromide (CTAB) method of Perez et al. (2012), then dissolved in TE buffer. DNA samples were quality-checked through electrophoresis on 1% agarose gel. DNA stocks of seven *O. sativa* cultivars from the Philippine Rice Research Institute (PhilRice) were added for comparison (Table 1).

**Table 1.** Summary information on the seven *O. sativa* varieties used in the study.

| Cultivar | Country of Origin | Remarks |
| --- | --- | --- |
| Nipponbare | Japan | Traditional japonica variety; reference sequence for *Oryza* sp. (Matsumoto et al., 2016) |
| NSIC Rc192 | Philippines | Inbred indica variety for rainfed lowland areas (Pinoy Rice Knowledge Bank, n.d.) |
| PSB Rc18 | Philippines | Inbred indica variety for irrigated lowland areas (Pinoy Rice Knowledge Bank, n.d.) |
| Li-Jiang-Xin-Tuan<br>-Hei-Gu | China | Cold-tolerant japonica rice variety (Pan et al., 2011) |
| Azucena | Philippines | Traditional upland variety; donor of the aroma gene, *frg* (Lorieux et al., 1996) |
| PSB Rc14 | Philippines | Inbred indica variety for rainfed lowland areas (Pinoy Rice Knowledge Bank, n.d.) |

**Figure 1.**
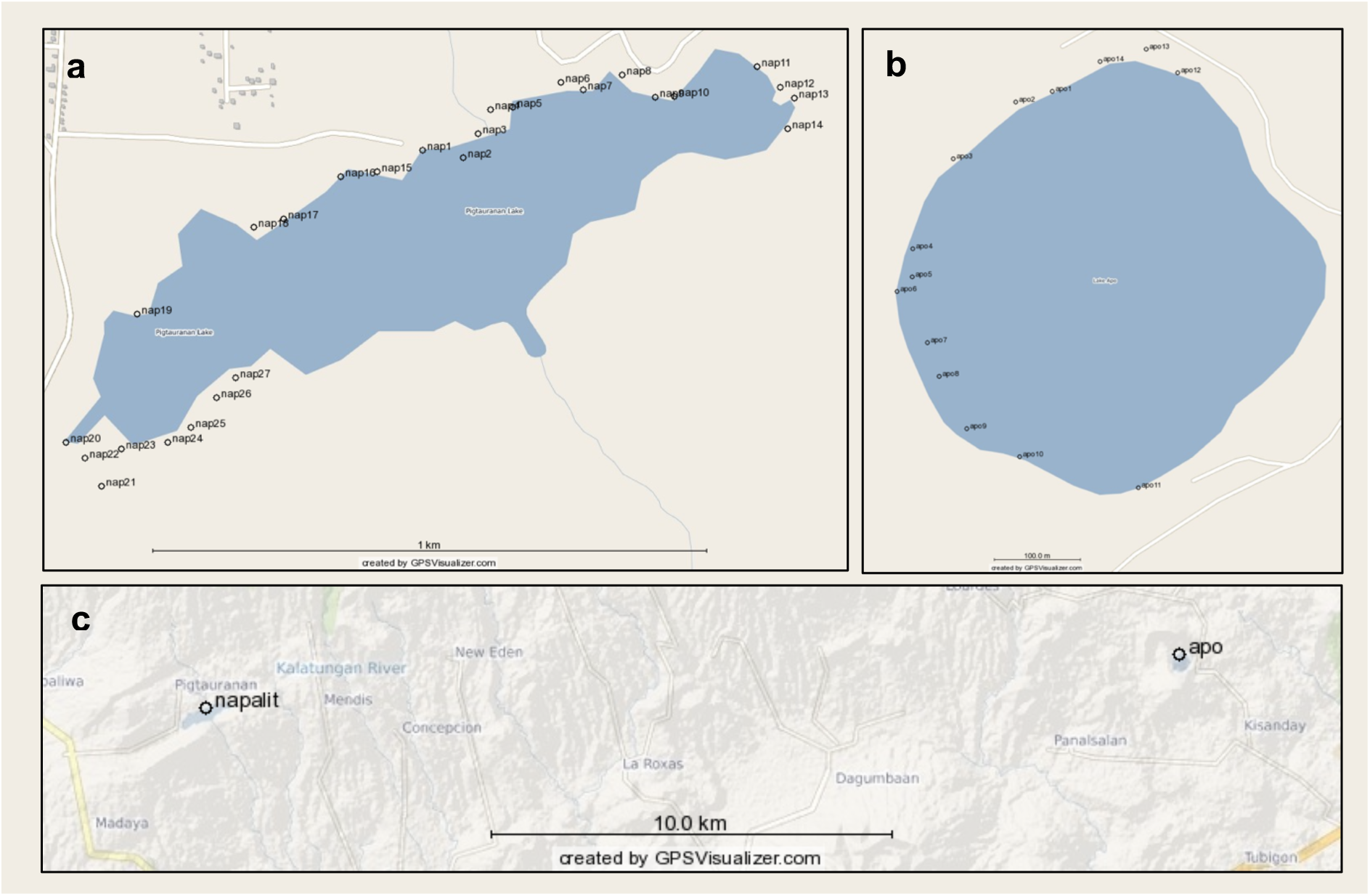
Sample locations of Rufi in (a) Lake Napalit and (b) Lake Apo. Distance between samples per lake was at least 30 m. (c) Distance between Lake Napalit and Lake Apo. Maps were generated using GPSVisualizer.com

One-hundred twenty-eight polymorphic genome-wide SSR markers for cultivated rice were initially selected, based on a 3-mb bin system to allow uniform marker distribution in the chromosomes, to evaluate the genetic variation among samples including the DNA stocks of seven *O. sativa* cv. (Figure 2). Polymerase Chain Reaction (PCR) was conducted per 6 μl reaction solution composed of 1.55 μl sdH20, 1.5 μl 10x PCR buffer, 0.6 μl 25 mM MgCl2, 0.35 μl 5mM dNTPs, 0.5 μl forward primer, 0.5 μl reverse primer, 1.0 μl 1:100 (concentrated suspension: storage buffer) *Taq* DNA polymerase, and 1.5 μl diluted DNA.

**Figure 2.**
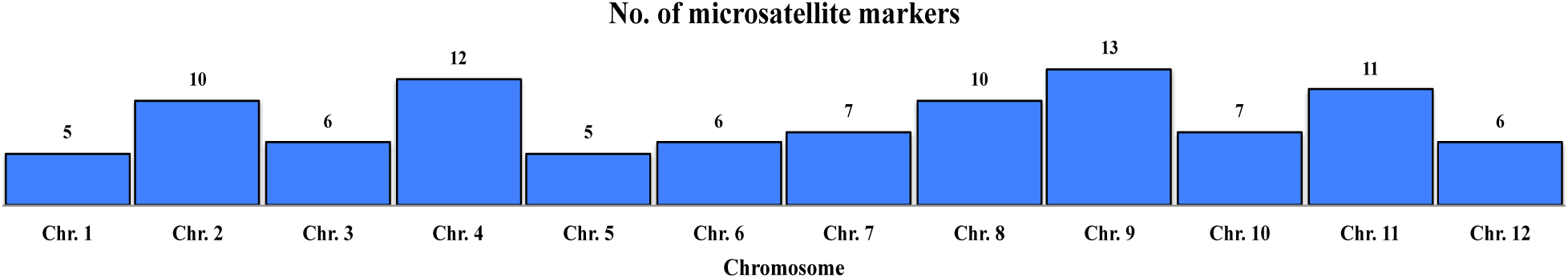
Distribution of the 98 polymorphic SSR loci across the 12 chromosomes selected based on a 3mb bin system of rice (*Oryza sativa* L.).

Each locus was amplified using a PTC-100® Thermal Cycler (Bio-Rad Laboratories, Inc., Hercules, CA, USA) for 2 hours and 30 min, following the PCR profile: 5 min initial denaturation at 94° C, followed by 35 cycles of 1 min denaturation at 94° C, 1 min annealing at 58° C, and 2 min extension at 72° C; then a final extension at 72° C for 5 min. PCR products were subjected to 6% polyacrylamide gel electrophoresis (PAGE) in 1x TBE buffer at 90 V for 1 to 2 hours depending on size of the PCR products. Gels were stained using ethidium bromide solution for 10-15 minutes. DNA bands were visualized using AlphaImager® HP system (ProteinSimple, San Jose, CA, USA). Detected allelic bands were scored from the fastest to the slowest migrating alleles by letters *a, b, c*, and so on.

### Data Analysis

A total of 128 SSR markers were initially selected to assess the genetic diversity and population structure of 41 Rufi samples along with 7 cultivated rice accessions for comparison which represents the different *O. sativa* ecotypes and subspecies, i.e. *indica* and *japonica* (Table 1). SSR markers with >20% missing band across all samples were excluded and only 98 markers were retained for further analysis (Figure 2). Rufi samples per lake were examined for genetic diversity per population. Seven rice cultivars added for comparison were grouped as the third population.

Genetic variation per population was assessed by calculating the percentage of polymorphic loci, mean number of alleles per locus, observed heterozygosity, expected heterozygosity, and Shannon information index. The Fixation index, Fst, (Weir and Cockerham, 1984) and analysis of molecular variance (AMOVA) were used to determine the degree of genetic divergence among populations, as well as, the primary source of genetic variation. The following scale for genetic differentiation was used: Fst < 0.05 (little), Fst = 0.05 – 0.15 (moderate), Fst = 0.15 – 0.25 (strong), and Fst > 0.25 (robust) (Hartl and Clark, 1997). Calculations were performed using GenAlEx ver. 6.5 (Peakall and Smouse, 2012). Moreover, population structure of Rufi and rice populations were evaluated by phylogenetic analysis where Nei’s genetic distance (Nei, 1972) was used to construct the pairwise genetic distance matrix for all 48 genotypes and neighbor-joining (NJ) tree was used for cluster analysis. The analysis was performed using PowerMarker ver.3.25 (Liu and Muse, 2004). MEGA 7: Molecular Evolutionary Genetics Analysis version 7.0 for bigger datasets (Kumar, et al., 2015) was used to construct and visualize the phylogenetic dendrogram.

To assess the patterns of genetic structure among Rufi and rice populations, a systemic Bayesian clustering approach applying Markov chain Monte Carlo (MCMC) estimation was performed using STRUCTURE ver. 2.3.2 (Pritchard et al., 2000), where the analysis was set using the admixture ancestry model, with number of assumed populations (K) set from 1 to 9, and each was ran 10x. Each run started with 200,000 burn-in periods followed by 200,000 MCMC iterations. MCMC was ran using a correlated allele frequency model based on the default frequency model information under the advanced option of STRUCTURE. Results from STRUCTURE were collated and imported to the web-based application Structure Harvester (Earl and vonHoldt, 2012) to calculate for Evanno’s delta K (Evanno, et al., 2005) which determined the optimal number of genetic groups.

## RESULTS

### Rufi populations showed low genetic diversity based on 98 SSR loci

Percentage of polymorphic loci (%P), mean number of alleles per locus (Na), mean number of effective alleles (Ne), observed heterozygosity (Ho), expected heterozygosity (He), and Shannon Information Index (I) were calculated to describe the genetic diversity of Rufi populations in comparison to *O. sativa* accessions. Results suggested that Napalit populations of Rufi were more genetically diverse than Apo populations (Table 2). However, genetic diversity of Rufi in both populations was lower compared to the cultivated rice accessions (Table 2). This higher diversity observed among *O. sativa* compared to Rufi might be due to its diverse countries of origin, and which were also bred for different ecotypes, and were also belong to different *O. sativa* subspecies (Table 1). Unlike the Philippine Rufi samples that were geographically isolated, with restricted gene flow, and that might have been predominantly reproducing asexually. In addition, the level of genetic diversity of these Rufi populations is lower compared to those reported by Zhou et al. (2003) in China (N_a_=10.6 among 10 SSR loci, 237 samples), Prathepha (2012) in Northeastern Thailand and Laos (N_a_=11.8571 among 7 SSR loci, 94 samples), and Ngu et al. (2010) in Malaysia (N_a_=14.8 alleles among 30 loci, 176 samples) despite using more SSR loci, as they had larger sample sizes which also came from several distant locations between sampled populations.

**Table 2.**
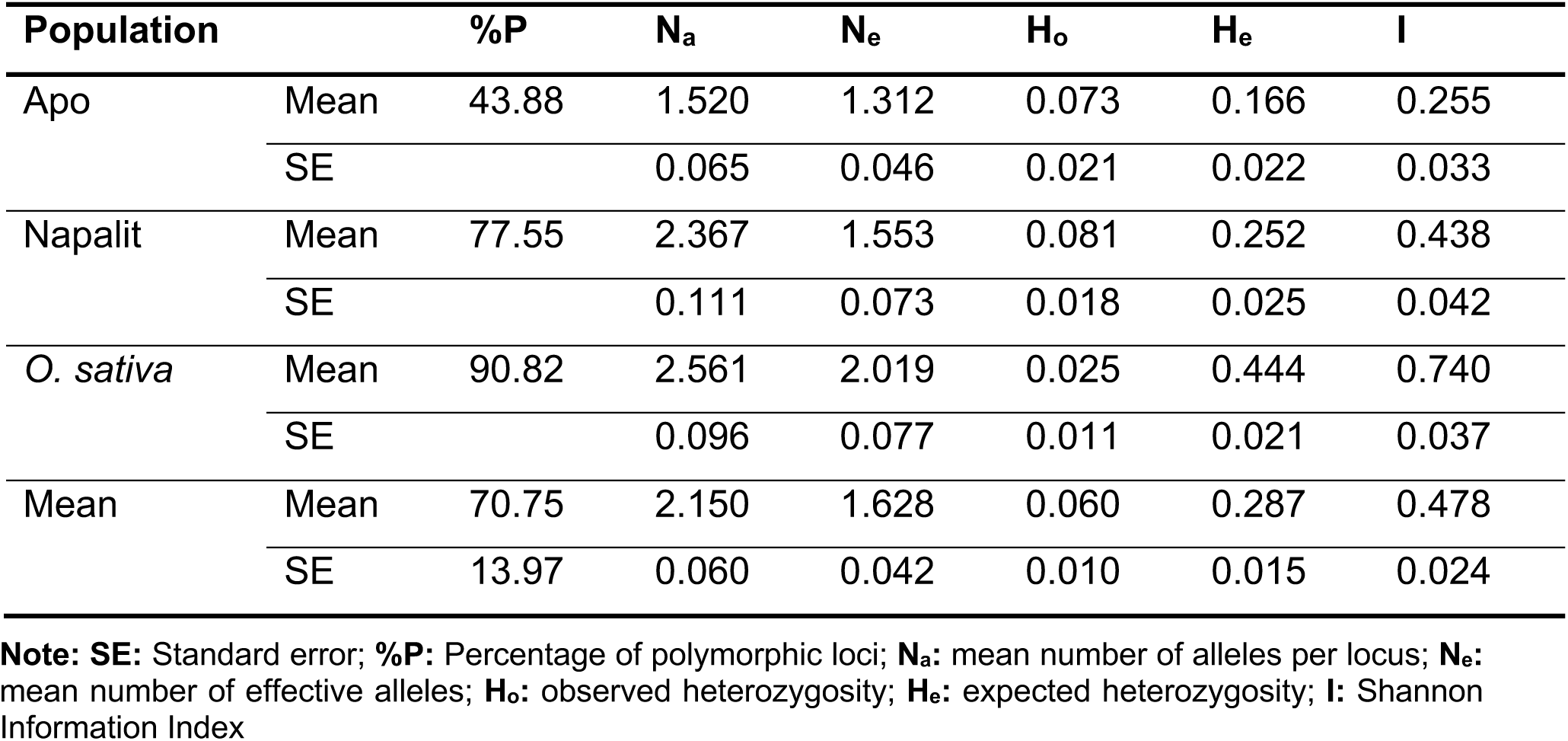
Genetic diversity parameters which showed that Rufi populations have lower diversity compared to the *O. sativa* cultivars.

| Population |  | %P | N<sub>a</sub> | N<sub>e</sub> | H<sub>o</sub> | H<sub>e</sub> | I |
| --- | --- | --- | --- | --- | --- | --- | --- |
| Apo | Mean | 43.88 | 1.520 | 1.312 | 0.073 | 0.166 | 0.255 |
|  | SE |  | 0.065 | 0.046 | 0.021 | 0.022 | 0.033 |
| Napalit | Mean | 77.55 | 2.367 | 1.553 | 0.081 | 0.252 | 0.438 |
|  | SE |  | 0.111 | 0.073 | 0.018 | 0.025 | 0.042 |
| *O. sativa* | Mean | 90.82 | 2.561 | 2.019 | 0.025 | 0.444 | 0.740 |
|  | SE |  | 0.096 | 0.077 | 0.011 | 0.021 | 0.037 |
| Mean | Mean | 70.75 | 2.150 | 1.628 | 0.060 | 0.287 | 0.478 |
|  | SE | 13.97 | 0.060 | 0.042 | 0.010 | 0.015 | 0.024 |

**Note:** **SE:** Standard error; **%P:** Percentage of polymorphic loci; **N<sub>a</sub>:** mean number of alleles per locus; **N<sub>e</sub>:** mean number of effective alleles; **H<sub>o</sub>:** observed heterozygosity; **H<sub>e</sub>:** expected heterozygosity; **I:** Shannon Information Index

Further, using the scale developed by Jamago and Cortes (2012) for Standardized Shannon-Weaver Diversity Index (H’ = 0.00 to 1.00), the degree of genetic diversity revealed by Shannon Information Index for both Apo and Napalit populations is poor (H’ = 0.01 – 0.45). This diversity level coupled with an equally low average number of alleles per locus per population might indicate that both Rufi populations in the Philippines are under severe genetic erosion and/or bottleneck.

### Apo and Napalit Rufi populations were distinct from each other

Population genetic structure was analyzed using the NJ tree based on Nei’s genetic distance and Bayesian clustering following the admixture ancestry model. Unrooted NJ tree showed clear separation between two *Oryza* species, where both Rufi populations were genetically closer to each other than to the rice cultivars (Figure 3a). Furthermore, Bayesian clustering results showed a major and minor peak of Evanno’s Δ*K* at *K = 2* and *K* = *3*, respectively (Figure 3b). At *K=2*, individuals were grouped according to species, i.e., *O. rufipogon* (green) and *O. sativa* L. (red) (Figure 3c). Also, two admixed samples (individuals) among Rufi populations were detected. At *K=3*, a new inferred cluster (blue) was revealed that comprised of all Napalit samples (Figure 3c). Six admixed individuals were detected in Napalit and Apo populations. When combined with the NJ tree results, Bayesian clustering indicated a robust genetic structuring of all Rufi individuals, especially at *K=3*. Results also showed that genetic clustering of Rufi individuals could be attributed to their geographic distribution. Fst and AMOVA determined the population genetic differentiation of Rufi individuals and rice accessions (Table 3). Results suggested that there is significant and robust population genetic differentiation (Fst=0.447, *p*=0.001) among Rufi populations from the *O. sativa* accessions. AMOVA results also indicated that variation within populations contributed significantly (55%, *P*=0.001) to this differentiation. Also, differences among populations had contributed significantly (45%, *p*=0.001) to population divergence.

**Table 3.** AMOVA summary table which showed that variation among and within populations is a significant contributor to population divergence.

| Source | Df | SS | MS | Est. Var. | %Var | P-value |
| --- | --- | --- | --- | --- | --- | --- |
| Among populations | 2 | 674.459 | 337.230 | 11.650 | 45% | 0.001 |
| Within populations | 93 | 1340.603 | 14.415 | 14.415 | 55% | 0.001 |
| Total | 95 | 2015.063 |  | 26.066 | 100 |  |

**Note:** **Df:** degrees of freedom; **SS:** sum of squares; **MS:** mean squares; **Est. Var.:** estimated variance; **%Var:** percentage of variance

**Figure 3.**
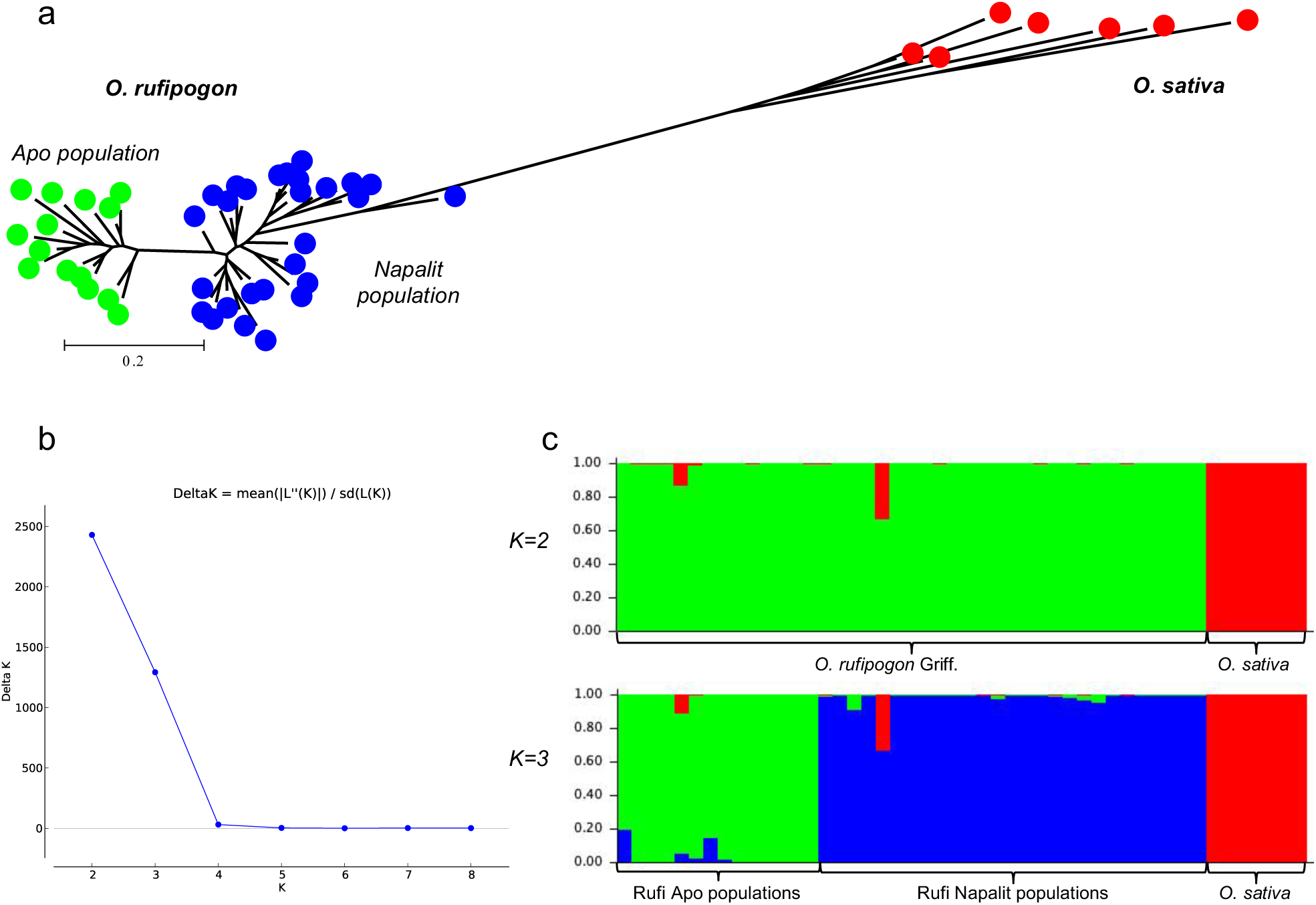
(a) Unrooted neighbor-joining tree based on Nei’s genetic distance which showed clear separation between the two *Oryza* species (b) estimated number of optimal clusters (K) calculated using the Evanno method which showed a major peak at K = 2, and a minor peak at K = 3, and (c) Bayesian clustering approach applying MCMC following the admixture ancestry model which showed that at K = 2 clustering is based on species and at K = 3 clustering is mainly based on the defined population groups.

## DISCUSSION

This is the first study to report on the molecular genetic diversity conducted on the known Rufi populations in the Philippines. Given their small population size especially for Apo, human settlement near the lakes and unsustainable farming practices such as slash and/or burn and overgrazing on the lakebanks based on the researchers’ silent monitoring since 2000, lack of awareness of nearby communities on Rufi’s importance for breeding programs based on informal conversations with locals, and influx of tourists in both lakes might be significant contributors to its low genetic diversity. Such might also indicate of a genetic bottleneck among Rufi populations in the country. Gao (2004) cited his unpublished data in 1996 and 1998 of field surveys over six years that identified anthropogenic factors had caused the extinction of 62.2% of *O. rufipogon* populations in China. Similarly, if the national and local governments do not adapt conservation measures and draw sustainability plans for Lakes Apo and Napalit, Rufi populations could eventually become locally extinct in the country.

Differentiation between Rufi and *O. sativa* indicates that domestication and plant breeding made these genetic resources genetically distant. Also, at some point of its development, an introgression event might have taken place between Rufi and *O. sativa* as per presence of admixed individuals in the two Rufi populations. Gliddon et al. (1987) suggested that gene flow in plants is a combination of pollen dispersal, seed dispersal, and clonal distribution. Rufi has a mixed breeding system, i.e. sexual and asexual. However, it is predominantly wind-pollinated and pollen dispersal is its primary mode of gene flow (Oka and Morishima, 1967). Due to a great distance that separates both lakes, approximately 24.47 km if measured in a straight line (Google, 2019), gene flow through pollen movement is improbable for the Rufi populations. Also, mature Rufi seeds have prominent long awns and are not suited for long-distance migration on their own, unless some are carried down through lake outlets if such exist. Seeds easily shatter and fall off from the mother plant. However, seeds with awns, barbs, and other similar structures can easily attach to passing animals while still on the plant rather than on the ground (Cruden, 1966). At Lake Napalit, water buffalos owned by nearby farmers often pass through the Rufi clumps. Seed dispersal can be also facilitated by small animals such as birds and rodents (Ngu et al., 2010). Gao (2004) proposed that clonal distribution, i.e. downstream movement of live ratoons and culms, is the leading mechanism for range expansion for Rufi. In the Philippines, movement of vegetative propagules is impossible since there is no known direct connection between the two lakes. However, there is popular folklore among locals living near Lake Napalit that the two lakes are connected, probably, by an underground river. Nonetheless, there is no current evidence that supports this folklore.

For this case, genetic differentiation between the Rufi populations can be attributed to their adaptation to unique geographic locations and climatic conditions. Lake Apo is situated at 640 masl whereas, Lake Napalit sits higher at 1,041 masl (Index Mundi, 2006). Balos (2013) and Balos and Jamago (2013) reported that Lake Napalit has higher rainfall (73.50 mm) and is relatively colder (23.5° C) than Lake Apo that recorded 26.33 mm and 25.3° C average rainfall and temperature, respectively. Differences in elevation and distance acted as geographic barriers which effectively restricted any form of gene flow. All these may have collectively contributed to population genetic structuring of the two natural Rufi populations. Further, habitat destruction due to unsustainable farming practices, human habitation along the lakebanks, and tourism have significantly reduced the Rufi populations in both lakes. As a result, individuals within each lake have been fragmented resulting to continuous inbreeding and clonal growth which magnify the genetic differentiation among and within populations over several generations (Zhou et al., 2003; Ngu et al., 2010).

## CONCLUSION

Lake Napalit Rufi populations are more diverse than the Lake Apo populations. However, Philippine Rufi has lower genetic diversity compared to other Rufi populations in China, Thailand, Laos, and Malaysia. Lake Apo and Lake Napalit populations are genetically distinct owing to the unique climate, significant elevation differences, and microenvironment conditions per lake. But this poor diversity is also possibly due to anthropogenic factors. Results indicate that the natural *O. rufipogon* populations on both lakes are in severe genetic bottleneck. If undesirable human activities are left uncheck or unregulated, Rufi in the Philippines may eventually become locally extinct. Given this grave threat and the importance of Rufi as raw material for rice breeding, urgent conservation plans should be in place in both lakes. With information on its population genetics already available, a sound decision on its effective and efficient conservation, sustainable use may be readily devised.

## ACKNOWLEDGMENTS

The authors would like to thank the local government units of Brgy. Guinoyoran, Valencia City and Brgy. Pigtauranan, Pangantucan, Bukidnon for letting us establish and conduct the study in Lake Apo and Lake Napalit, respectively. They would also like to express their gratitude to the Philippine Rice Research Institute-Central Experiment Station for supporting and funding this research project.

## REFERENCES

Balos, J.L. (2013). Comparative Morpho–Ecological Characterization of *In Situ* Populations of *Oryza rufipogon* Griff. in Lakes Apo and Napalit, Bukidnon. Unpublished Undergraduate Thesis. Central Mindanao University, Musuan, Bukidnon. (Available at the CMU Library)

Balos, J.L. and Jamago, J.M. (2013). Comparative characterization of *in situ Oryza rufipogon* Griff. populations in Lakes Apo and Napalit, Bukidnon. Transactions of the National Academy of Science and Technology, 35(1):14 from https://bit.ly/2BbUOpz

Bon, S.G., and Borromeo, T.H. (2003). Discovery and re–discovery of wild rice populations in the Philippines. Philippine Journal of Crop Science, 27: 53–58.

Bui, C., and Nguyen, T. (2017). QTL analysis on rice genotypes adapted to acid sulfate soils in the Mekong river delta, Vietnam. Vietnam Journal of Science, Technology and Engineering, 59(4), 26–31.

Cruden, R.W. (1966). Birds as agents of long-distance dispersal for distinct plant groups of the temperate western hemisphere. Evolution, 20: 517–532.

Earl, D.A., and Von Holdt, B.M. (2012). STRUCTURE HARVESTER: a website and program for visualizing STRUCTURE output and implementing the Evanno method. Conservation Genetics Resources, 4(2), 359–361 doi: 10.1007/s12686-011-9548-7.

Evanno, G., Regnaut, S., and Goudet, J. (2005). Detecting the number of clusters of individuals using the software structure: a simulation study. Mol. Ecol. 14: 2611–2620. doi: 10.1111/j.1365-29X.2005.02553.x.

Gao, L.Z., Schaal, B.A., Zhang, C.H., Jia, J.Z., and Dong, Y-S. (2002). Assessment of population genetic structure in common wild rice *Oryza rufipogon* Griff. using microsatellites and allozyme markers. Theor Appl Genet., 106: 173–180.

Gao, L.Z. (2004). Population structure and conservation genetics of wild rice *Oryza rufipogon* (Poaceae): a region-wide perspective from microsatellite variation. Mol Ecol., 13:1009–1024

Gliddon, C., Belhassen, E., and Gouyon, P.H. (1987). Genetic neighborhoods in plants with diverse systems of mating and different patterns of growth. Heredity, 59: 29–32.

GOOGLE. (2019). Bukidnon. Retrieved from http://tiny.cc/zjtj6y on 11 July 2019.

INDEX Mundi. (2006). Philippines. Retrieved from http://tiny.cc/xktj6y on 11 July 2019.

Jamago, J.M., J.L. Balos, M.L. Fuentes, J.D. Genilla, A.M. Racines, J.M. Remollo, and M.A. Tejada. (2012). Wild Rice in a Forgotten Paradise. Plant Genetic Resources Exhibit Poster Presentation. Second semester, 2011-2012. Department of Agronomy and Plant Breeding, College of Agriculture, CMU, Musuan, 8710 Bukidnon.

Jamago, J.M. and Cortes, R.V. (2012). Seed diversity and utilization of the upland rice landraces and traditional varieties from selected areas in Bukidnon, Philippines. IAMURE Int. J. Ecol. Conserv., 4: 112–130.

Kumar, S., Stecher G., Li M., Knyaz, C., and Tamura, K. (2018). MEGA X: Molecular volutionary genetics analysis across computing platforms. Molecular Biology and Evolution, 35: 1547–1549.

Li, R.B., Li, L.S., Wei, S.M., Wei, Y.P., Chen, Y.Z., Bai, D.L., Yang, L., Huang, F.K., Lv, W.L., Zhang, X.J., Li, X.Y., Yang, X.Q., and Wei, Y.W. (2010). The evaluation and utilization of new genes for brown planthopper resistance in common wild rice (*Oryza rufipogon* Griff.). Mol Entomol., 4: 365–371.

Lorieux M, Petrov M, Huang N, Guiderdoni E, Ghesquière A (1996) Aroma in rice: genetic analysis of a quantitative trait. Theor App Genet 93:1145–1151

Matsumoto, T., Wu, J., Itoh, T., Numa, H., Antonio, B., & Sasaki, T. (2016). The Nipponbare genome and the next-generation of rice genomics research in Japan. In Rice (Vol. 9, Issue 1). Springer New York LLC. https://doi.org/10.1186/s12284-016-0107-4

Nei, M. (1972). Genetic distance between populations. American Naturalist, 106: 283–392.

Ngu, M.S., Sabu, K.K., Lim, L.S., Abdullah, M.Z., and Wickneswari, R. (2010). Genetic structure of *Oryza rufipogon* Griff. natural populations in Malaysia: implications for conservation and genetic introgression of cultivated rice. Tropical Plant Biol., 3: 227–239.

Nguyen, B.D., Brar, D.S., Bui, B.C., Nguyen, T.V., Pham, L.N., and Nguyen, H.T. (2003). Identification and mapping of the QTL for aluminum tolerance introgressed from the New source, *Oryza rufipogon* Griff., into indica rice (*Oryza sativa* L.). Theor Appl Genet., 106: 583–593.

Oka, H.I., and Morishima, H. (1967). Variations in the breeding systems of a wild rice, *Oryza perennis*. Evolution, 21: 249–258.

Orloci, L. (1978). Multivariate analysis in vegetation research. The Hague: Dr W. Junk B.V.

Pan, Y. J., Wang, W. S., Zhao, X. Q., Zhu, L., Fu, B., & Li, Z. (2011). DNA methylation alterations of r ice in response to cold stress. Plant Omics, 4(7), 364–369.

Peakall, R. and Smouse, P.E. (2012). GenAlEx 6.5: genetic analysis in Excel. Population genetic software for teaching and research-an update. Bioinformatics, 28: 2537–2539.

Perez, L.M., Domingo P.H., Tabanao, D.A., and Manigbas, N.L. (2012). DNA fingerprinting in hybrid rice: its application in varietal purity testing. Philippine Rice Research Institute, Science City of Muñoz, Nueva Ecija, Philippines.

PINOY RICE KNOWLEDGE BANK. (n.d.). Rice Varieties. Retrieved from https://www.pinoyrice.com/rice-varieties/ on 5 July 2020.

Pritchard, T., Jamjod, S., and Donnelly, P. (2000). Inference of population structure using multilocus genotype data. Genetics, 155: 945–959.

Prathepha, P. (2012). Genetic diversity and population structure of wild rice, *Oryza rufipogon* from Northeastern Thailand and Laos. Australian Journal of Crop Science, 6(4): 717–723.

Shibata, Y., Cabunagan, R.C., Cabauatan, P.Q., and Il-Ryong, C. (2007). Characterization of *Oryza rufipogon*-derived resistance to tungro disease in rice. Plant Disease - PLANT DIS, 91: 1386-1391. 10.1094/PDIS-91-11-1386.

Song, Z.P., Li, B., Chen, J.K., and Lu, B.R. (2005). Genetic diversity and conservation of common wild rice (*Oryza rufipogon*) in China. Plant Species Biol., 20: 83–92.

Weir, B.S. and Cockerham, C.C. (1984). Estimating F-statistics for the analysis of population structure. Evolution, 38: 1358–1370.

Zhou, H.F., Xie, Z.W., and GE, S. (2003). Microsatellite analysis of genetic diversity and population genetic structure of a wild rice (*Oryza rufipogon* Griff.) in China. Theor Appl Genet., 107: 332–339.

